# InterPepRank: Assessment of Docked Peptide Conformations by a Deep Graph Network

**DOI:** 10.1101/2020.09.07.285957

**Authors:** Isak Johansson-Åkhe, Claudio Mirabello, Björn Wallner

## Abstract

**Motivation:** Peptide-protein interactions between a smaller or disordered peptide stretch and a folded receptor make up a large part of all protein-protein interactions. A common approach for modelling such interactions is to exhaustively sample the conformational space by fast-fourier-transform docking, and then refine a top percentage of decoys. Commonly, methods capable of ranking the decoys for selection in short enough time for larger scale studies rely on first-principle energy terms such as electrostatics, Van der Waals forces, or on pre-calculated statistical pairwise potentials.

**Results:** We present InterPepRank for peptide-protein complex scoring and ranking. InterPepRank is a machine-learning based method which encodes the structure of the complex as a graph; with physical pairwise interactions as edges and evolutionary and sequence features as nodes. The graph-network is trained to predict the LRMSD of decoys by using edge-conditioned graph convolutions on a large set of peptide-protein complex decoys. InterPepRank is tested on a massive independent test set with no targets sharing CATH annotation nor 30% sequence identity with any target in training or validation data. On this set, InterPepRank has a median AUC of 0.86 for finding coarse peptide-protein complexes with LRMSD*<*4Å. This is an improvement compared to other state-of-the-art ranking methods that have a median AUC of circa 0.69. When included as selection-method for selecting decoys for refinement in a previously established peptide docking pipeline, InterPepRank improves the number of Medium and High quality models produced by 80% and 40%, respectively.

**Availability:** The program is available from: *http://wallnerlab.org/InterPepRank*

**Contact:** Björn Wallner

**Supplementary information:** Supplementary data are available at *BioRxiv* online.

## 1 Introduction

Interactions between a short stretch of amino acid residues and a larger protein receptor, referred to as peptide-protein interactions, make up ap-proximately 15-40% of all inter-protein interactions (Petsalaki and Russell, 2008), and are involved in regulating vital biological processes (Midic *et al.*, 2009; Tu *et al.*, 2015). These short peptide stretches are often disordered when unbound or part of larger disordered regions (Petsalaki and Russell, 2008; Neduva, Victor *et al.*, 2005). Because of their inherent flexibility predicting peptide-protein interaction complexes is difficult.

However, since the peptide ligand is a smaller molecule, it is possible to exhaustively sample the binding space by Fast-Fourier Transform docking (FFT-docking). Classically, this requires a close to correct rigid receptor and ligand, but as shown in Alam *et al.* 2017, a set of poses derived from protein fragments with sequence similar to the peptide can consistently produce conformations near the native bound conformation. While the strength of FFT-docking is the exhaustive search of docking space, the problem, as we will show, is that the energy function is a limited approximation of the binding affinity, and thus even though the method samples many near-native decoys it often fails to separate them from poor decoys. Additionally, rigid-body docking needs to be followed up with much more computationally expensive refinement to reliably produce native contacts and binding modes.

A workflow previously shown to be successful is to rescore and refine promising decoys using a more advanced energy-function, such as Rosetta (Raveh *et al.*, 2010). Indeed, previous works have shown great success in the peptide-protein docking area by combining FFT-based docking with Rosetta refinement (Alam *et al.*, 2017). However, because of the computational cost in running refinement, only a tiny subset if the FFT-generated decoys can be used. This selection is based on the energy function of the FFT method, resulting in unnecessary refinement of several decoys unsalvageable by the refinement protocol.

An improvement to this approach would be to run all decoys through a fast and accurate re-scoring algorithm to select decoys for refinement, rather than relying on energy functions constrained to FFT-compatibility. Many methods have been developed for the rescoring of protein-protein complexes, some examples include: PyDock evaluates decoys based on pairwise electrostatic potentials and desolvation energy (Pallara *et al.*, 2017; Cheng *et al.*, 2007). Zrank and Zrank2 utilizes van der Waals in excess of electrostatics and desolvation (Pierce and Weng, 2007, 2008). OPUS-PSP uses orientation-dependent packing and knowledge-based repulsive energy (Lu *et al.*, 2008). SIPPER uses statistical residue-pair potentials derived from a curated interaction-set (Pons *et al.*, 2011). However, most rescoring methods for docked complexes are not optimized for the peptide-protein problem, and to the best of our knowledge still rely on either physics-based manually defined energy-functions, or knowledge-based and empirically tested energy-function approximations, that do not take evolutionary information of the target into account. More advanced methods using machine learning, like ProQDock (Basu and Wallner, 2016), have running times unfeasible for application on a large set of decoys and are better suited for evaluating a small set of refined models.

Within structural bioinformatics, Graph Convolutional Networks (GCNs) have seen increased use recently through applications such as PipGCN which uses a pairwise classification architecture and evolutional features to predict protein-protein binding sites (Fout *et al.*, 2017), the GCN of (Gligorijevic *et al.*, 2019) which feeds pre-trained LSTM sequence feature extraction to a graph network to classify protein function, EGCN which uses edge-based graph convolutions to score protein-protein complexes with the use of simple features such as side-chain charge and hydrophobicity (Cao and Shen, 2019), and GCNN of (Zamora-Resendiz and Crivelli, 2019) which encodes spatial information into the residue-nodes to classify proteins. A Graph Convolutional Network eschews the spatial limitations of a classic convolutional network and allows for the direct definition of spatial relationships on a case-by-case basis. It passes information along pre-defined edges between nodes rather than by proximity in the input-matrix, while still allowing for complex information, such as evolutionary information, to be encoded in the nodes.

In this work, we present a fast, novel, rescoring algorithm utilizing GCNs to represent the peptide-protein complex with the added context of evolutionary information. The GCNs is capable of quickly sift through and rank the complete space of conformations generated by FFT-methods and improving the selection of decoys for subsequent refinement.

## 2 Materials and Methods

### 2.1 IPD0220 Dataset

A set of 6,857 interacting peptide-protein pairs taken from the PDB (Berman *et al.*, 2000) at 15/10/2018 was redundancy reduced by 30% sequence identity down to 687 pairs. In this case, a peptide-protein interaction is defined as a peptide of 25 or fewer residues sharing a contact surface of at least 400Å^2^ with a receptor of 50 residues or more. The full dataset of 687 pairs was further randomly divided into partial sets of circa 50 pairs each, with the requirement that no receptor in any of these sets could share a CATH superfamily annotation (Dawson *et al.*, 2017) with any receptor from any other of the sets. In the case a receptor lacked CATH annotation, it was aligned to all other receptors with TM-align (Zhang and Skolnick, 2005) and classified as similar if it had a TM-score*>*0.4.

One of the sets was randomly selected as a validation set. The remaining 13 sets were all used for testing in a jack-knifing-like scheme with a unique training set generated for each test set. The network parameters and architecture was optimized on the validation set once, while early stopping was performed with the help of the validation set for all tests.

The complete data set is published available here (Johansson-Åkhe *et al.*, 2020b)

#### 2.1.1 Test and Validation Set Generation

To achieve more variety and difficulty in the sets, peptide conformations were generated rather than using the native conformations alone. For each target of each of the test and validation sets, 50 different peptide conformations were generated using the Rosetta fragment-picker (Gront *et al.*, 2011). Each of these conformations were exhaustively docked on the surface of their receptors by PIPER using FFT-docking (Kozakov *et al.*, 2006), generating 70,000 decoys for each conformation for each target in the sets, in total close to 2.5 billion (70,000×50×687) decoys. To enable rapid method development, only a random subset 2,500 decoys per target were used in the validation.

To ensure that each of the validation and test sets represent the total dataset, the distribution of LRMSD of all decoys for each set were compared with Kolmogorov-Smirnov test to the distribution of all data not in the set. These tests showed that the sets could not be said to represent different distributions (P*>*0.37).

#### 2.1.2 Training Set Generation

A unique training set was constructed for each test set, to allow a broad variety in training decoys while simultaneously ensuring the method was never trained on examples too similar to what it was tested on. When testing on one of the test sets, each of the original non-redundancy reduced 6,587 peptide-protein pairs which did not share a CATH annotation nor 30% sequence identity with the test set in question nor the validation set were used for training. For each of these, peptide conformations were generated as for the test and validation sets, but 4,375 decoys for each conformation were used rather than 70,000. A balanced dataset was achieved by only permitting each target to contribute as many incorrect decoys as it could contribute correct decoys (definition of a correct decoy can be found in the Metrics section below). Additionally, to avoid bias, the number of decoys which was included in the finalized training sets by each CATH superfamily was limited by the median number of decoys each CATH superfamily could contribute, resulting in each peptide-protein pair contributing on average 426 decoys, divided equally between correct and incorrect decoys. This resulted in the number of decoys in each unique training set totaling on average 1,048,386.

#### 2.1.3 Expanded Analysis set

One of the test-sets was randomly selected as an Expanded Analysis set for running extra analysis and refinement on. This allows for the use of computationally slow refinement methods to test the final impact the methods in this paper can have on full docking protocols, and additionally allows for comparison to relatively slow re-scoring methods such as pyDock3.

### 2.2 Metrics

Several different metrics are used to evaluate the performance of InterPepRank and other methods benchmarked.

#### 2.2.1 Metrics for Rigid-Body-Docked decoys - Correct Decoys

In the perspective of ranking rigid-body-docked decoys, a decoy is defined as correct if it has the peptide positioned within 4.0Å ligand root-mean square deviation (LRMSD) of the native conformation. This limit was selected both as it is within the reported limit of when the Rosetta Flex-PepDock refinement protocol can reliably refine decoys to sub-Ångström precision (Raveh *et al.*, 2010), and since it is below the CAPRI limit for an acceptable prediction of a docked peptide-protein complex (Lensink *et al.*, 2017).

Since the goal of the method presented herein is not to produce final high-quality predictions, but rather to select which rigid-body-docked decoys are worth refining, a higher cutoff than the sub-Ångström high-quality model cutoff was selected.

#### 2.2.2 Metrics for Refined Models - Acceptable, Medium, High

Later in the paper, the quality of models refined from the rigid-body-docked decoys by the Rosetta FlexPepDock refinement is discussed. In this case, molecular details of interaction and sub-Ångström positioning of the peptide becomes relevant, and the more lax definition of a correct decoy established earlier needs to be supplemented. The CAPRI standard for assessing refined peptide-protein docked complexes can be found in Table 1.

**Table 1:** CAPRI criteria for peptide-protein docked model quality (Lensink *et al.*, 2017). iRMSD is the root mean square deviation of residues at the native interface. Fnat is the fraction native residue-residue contacts recalled. A model which is at least High quality will also be at least Medium and Acceptable quality, like a Medium quality model will also be at least Acceptable quality.

| *Model Quality* | LRMSD | iRMSD | fnat |
| --- | --- | --- | --- |
| Acceptable | < 5.0 Å | < 2.0 Å | > 0.2 |
| Medium | < 2.0 Å | < 1.0 Å | > 0.5 |
| High | < 1.0 Å | < 0.5 Å | > 0.8 |

#### 2.2.3 ROC

A receiving operand characteristic-curve (ROC-curve) measures how two metrics change in relation to each-other, here False Positive Rate (FPR) and True Positive Rate (TPR), as the threshold for scoring is varied.

False Positive Rate is defined as:

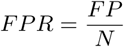

where *F P* is the number of decoys incorrectly identified as correct and *N* is the total number of incorrect decoys in the set.

True Positive Rate is defined as:

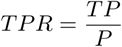

where *T P* is the number of correctly identified correct decoys and *P* is the total number of correct decoys in the set.

### 2.3 Representation and Architecture

#### 2.3.1 Decoy Representation

The graph network utilizes edge-conditioned graph convolutions, where the input is a set of nodes, each one described by a set of input features, as well as edges of different types between nodes. In this case, nodes represent individual residues of a decoy and edges denote different types of interactions between couples of residues. In this study, the node features used were Amino Acid Code (one-hot encoded, 21 values to account for unknown residues), a Position-Specific Scoring Matrix (21 values including gaps, Eq. 1), Self-Entropy (21 values, including gaps, Eq. 2), and one variable denoting if the residue belongs to the peptide or receptor (1 value), for a total of 64 features. Multiple sequence alignments for calculation of PSSM and self-entropy were acquired by running HHblits 2.0.15 with uniclust30 2016 03 as the database, running 2 iterations with maximum pairwise sequence identity of 90% and E-value inclusion threshold of 0.001.

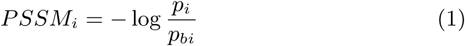

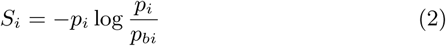

where *p*_*i*_ is the frequency of the amino acid on position *i* recurring at that position in the multiple sequence alignment and *p*_*bi*_ is the background probability of that amino acid.

Four types of one-hot-encoded edges were used: self-edges to allow the passing on of information to the same residue in consecutive layers, covalent edges denoting the existence of a covalent bond between two residues, proximity edges between each pair of residues with any heavy atoms within 4.5Å, and to speed up calculations through filtering where convolutions can be performed a summarizing identity edge feature set to 1 if any other edge feature was present. For each decoy, only 100 nodes and the vertices that connected them were used; the peptide and the residues of the receptor closest to the peptide. This means that if the peptide is 25 residues long and the receptor 170 residues, the 100 nodes would include the 25 peptide residues as well as the 75 residues from the receptor which were closest to any peptide amino acid. For cases where all the residues in the complex amount to fewer than 100, zero-padding is used.

#### 2.3.2 Target Function

To facilitate training, the raw LRMSD values were normalized to the [0,1] range using the same normalization scheme as in Levitt and Gerstein 1998:

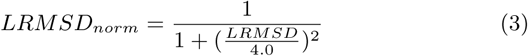

Since networks are often observed learning quicker from classification tasks, the problem was formulated as a classification problem by predicting to which bin of *LRMSD*_*norm*_ a decoy belonged to. The number of bins were varied from 2-4 evenly spread in the [0,1] range. The final predicted score, *S*, was a sum of the *LRMSD*_*norm*_ bins weighted by the predicted indeprobability:

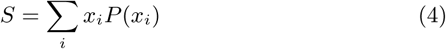

where *i* is the number of bins, *x*_*i*_ the center of bin *i* and *P* (*x*_*i*_) the predicted probability for bin *x*_*i*_.

#### 2.3.3 Network Architecture

The network was implemented as a feed-forward graph convolutional network, see Figure 1. Before any convolutions were applied, the node feature Amino Acid Code (one-hot encoded 21 value feature) was passed through an Embedding Layer to reduce dimensionality and thus also the number of weights, limiting overfitting. Embedding layers are small network architectures for mapping discrete labels onto continuous space, and are frequently used in areas such as language processing (Mikolov *et al.*, 2013), and have previously successfully been used to describe amino acids (Mirabello and Wallner, 2019).

**Figure 1:**
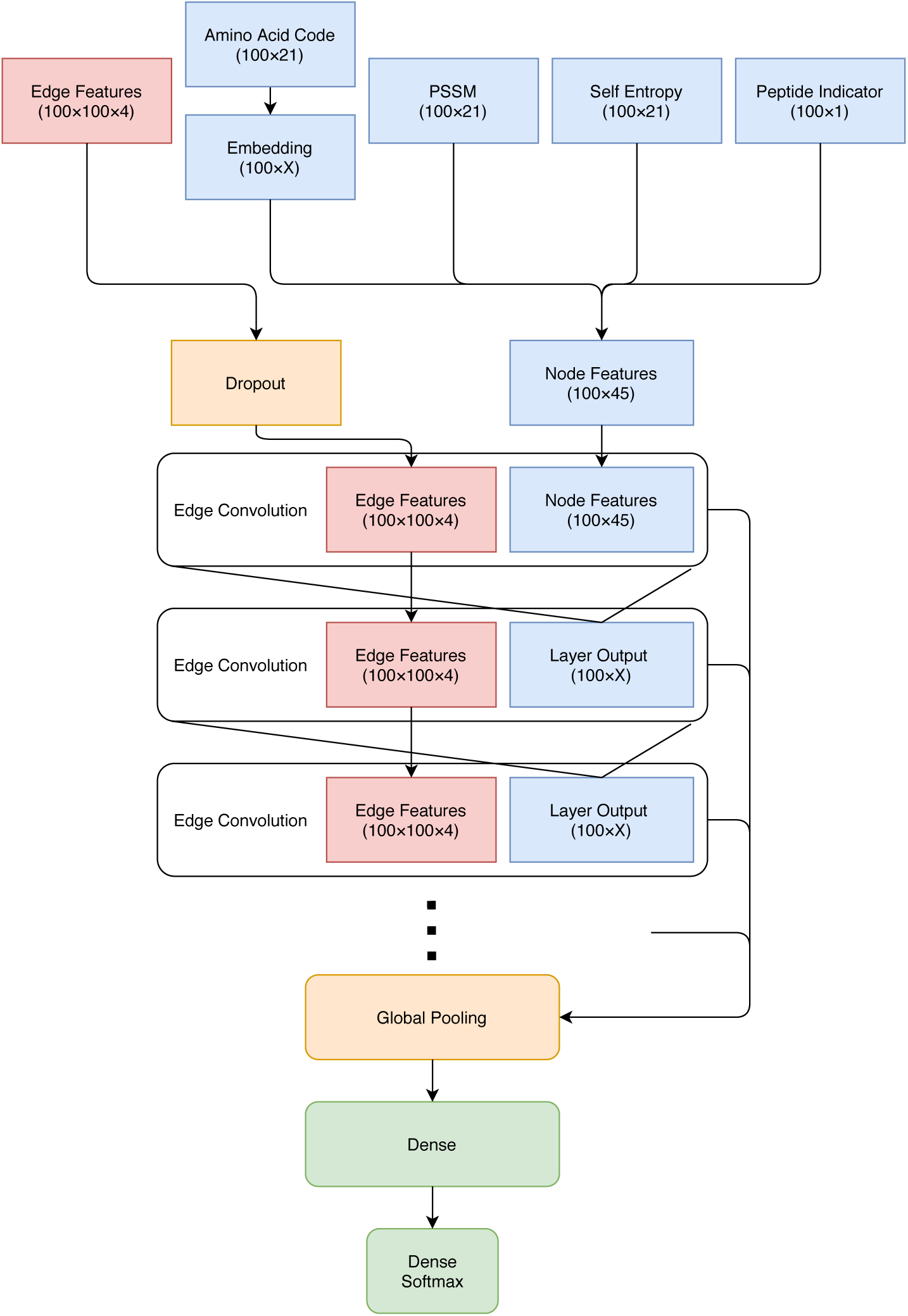
The basic architecture of the InterPepRank nets. Note that the output from the Edge Convolution and Embedding layers have been denoted as X, since different values were sampled. Throughout the net, the ReLU activation function is applied after each convolution.

Next, the node features were passed through a varied number of Edge Convolution Layers (Simonovsky and Komodakis, 2017), with ReLU activation between each layer, taking the output of the last layer as node features for the next, while keeping the same edge features throughout. Edge Convolution Layers learn filters as a function of both node and edge features, and apply these filters along the edges of the graph.

The output from each convolution layer was concatenated together before global pooling of all node features, followed by 2 dense layers before prediction. During training, a dropout ranging between 0.1 and 0.25 was applied to the edge features. Different sizes of the Edge Convolution and Dense outputs were explored, as well as different methods for Pooling, see below.

The network was implemented using Spektral and Keras (Chollet *et al.*, 2015), using the layers proposed by Simonovsky and Komodakis 2017, and trained with the TensorFlow backend (Abadi *et al.*, 2015).

#### 2.3.4 Parameter Optimization and Ensembling

Parameters and hyperparameters of the nets were optimized to maximize performance on the validation set. Additionally, to increase the predictive power of InterPepRank with little impact on run-time, some of the best-performing trained nets were ensembled by averaging their outputs.

## 3 Results and Discussion

In this work we have developed InterPepRank, a machine-learning based method which encodes the structure of peptide-protein complex as a graph; with physical pairwise interactions as edges and evolutionary features such as PSSM and sequence conservation as nodes. The graph representation is trained to predict the LRMSD of decoys by using edge-conditioned graph convolutions on a large set of peptide-protein complexes. Different network architectures were tried and the nine best (0-8) on validation are shown in Figure 2, details on specific network architectures in the Supplementary Information. To maximize performance a subset of these were used as an ensemble predictor, averaging over their output; best ensemble used all networks except 5 and 6.

**Figure 2:**
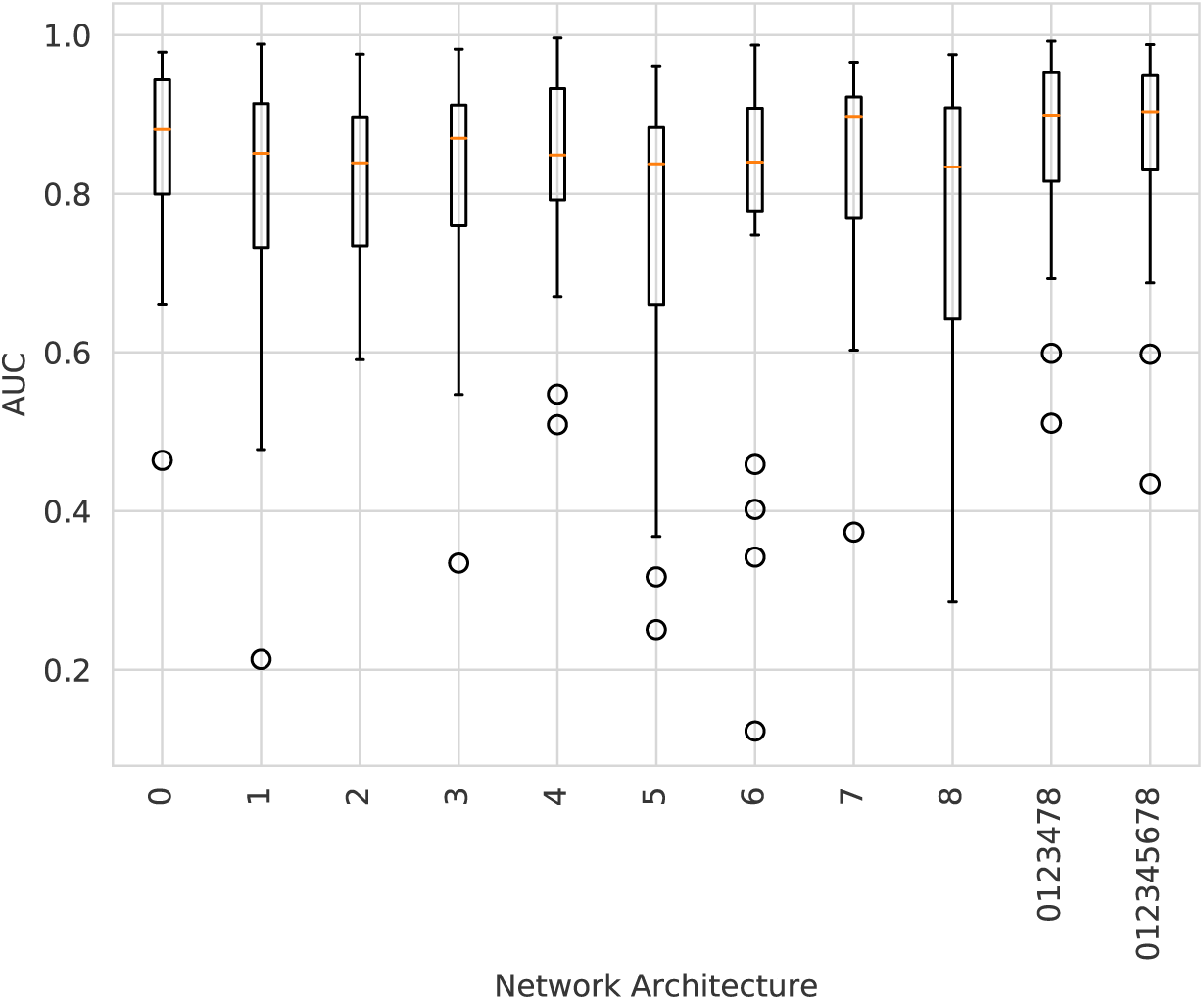
AUC on validation targets for the different individual network architectures, the final ensemble method and an ensemble including all architectures. The network architectures are numbered from 0 through 8, and the ensemble methods are named after the included architectures. The ensemble 0123478 shows optimal performance on the validation data. A detailed description of the architectures and their differences can be found in the Supplementary Information.

### 3.1 Comparison to Established methods

To put the performance of InterPepRank into perspective, we compared its ranking strength to several established state-of-the-art docked complex scoring methods: PIPER (Kozakov *et al.*, 2006), pyDock3 (Cheng *et al.*, 2007), and Zrank (Pierce and Weng, 2007). PIPER was run with the same rotation and energy matrices as when part of the peptide-protein docking protocol PIPER-FlexPepDock (Alam *et al.*, 2017). Although both pyDock3 and Zrank are optimized for the scoring of protein-protein complexes, pyDock3 was recently shown to efficiently identify near-native peptide decoys in the sixth CAPRI edition (Pallara *et al.*, 2017), and Zrank is included as one of the currently leading fast available protein-protein scoring functions according to (Moal *et al.*, 2013) and (Yan and Huang, 2019), amongst others. It should be noted that PIPER was also used to generate the decoys of this study.

As discussed in the introduction, currently there is a lack of ready-to-use scoring functions developed specifically for peptide-protein complexes. Technically, the Rosetta FlexPepDock protocol (Raveh *et al.*, 2010) can be run in a mode which only applies its scoring function without changing the structure. However, Rosetta utilizes a fine-grained scoring algorithm, and the structures needs to be minimized using the Rosetta relax protocol to be scored properly.

The primary objective of this study was to develop a method for selecting decoys for further refinement, which means the metric of interest is the ability of the method to rank decoys for individual targets. InterPepRank has a higher average AUC compared to both Zrank and PIPER, see Figure 3. The median AUC is 0.86 for InterPepRank compared to 0.65 and 0.69 for Zrank and PIPER, respectively. The same trend holds true against pyDock3 and Rosetta FlexPepDock scoring as well on the Expanded Analysis set, see Supplementary Figure 1.

**Figure 3:**
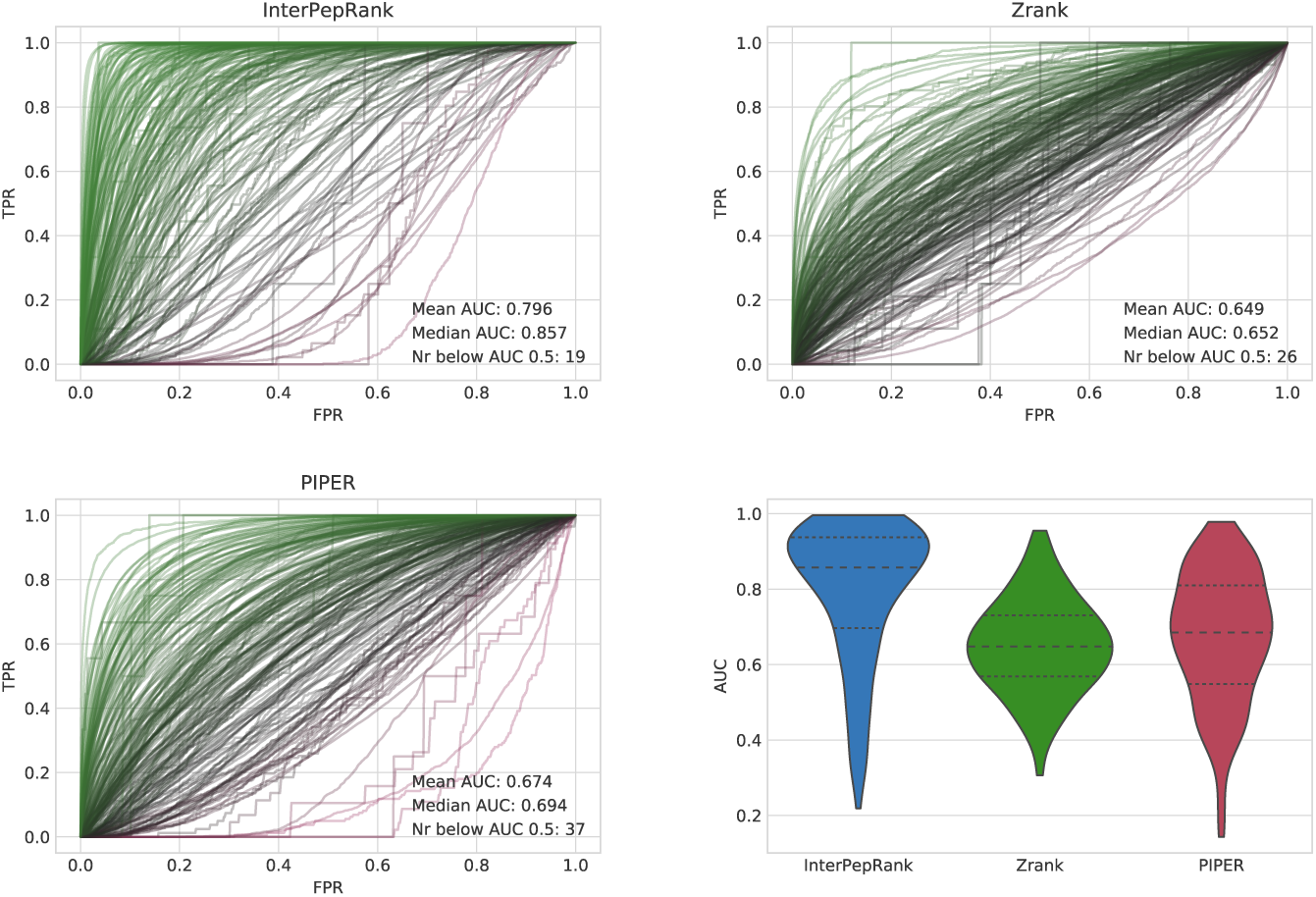
ROC-curves for the different methods, each target is represented by 1 curve, and a violin-plot over the distributions of AUCs. The area under the curve (AUC) displayed in the graphs is the average and median over all targets. Note that while the standard deviation of AUC is higher for InterPepRank, it still only has 19 targets with an AUC below 0.5, compared to 37 and 26 targets for PIPER and Zrank, respectively.

Another metric of interest is the ability to rank decoys even between targets, testing if the methods are capable of absolute decoy ranking. This was tested on the Expanded Analysis set. Overall InterPepRank is much better than Zrank, pyDock3, and PIPER at assessing the absolute quality of individual decoys in comparison to both other decoys from the same target and between targets, even when scores are normalized by receptor and peptide length, see Figure 4, with an AUC of 0.90 compared to 0.73 and 0.70 for pyDock3 and Zrank, respectively. PIPER is slightly worse with AUC 0.66. This is expected as the other scoring methods are dependent on complex size and primary structure composition. The overall AUCs of Figure 4 mirror the median AUCs of Figure 3 when PIPER and Zrank are normalized by receptor and ligand length. This indicates the methods tested should be roughly equally able to pick out refineable decoys from a pool of different receptors and ligands as they should be at ranking internally, implying the results from the tests focusing on internal ranking could be generalized to an overall selection case.

**Figure 4:**
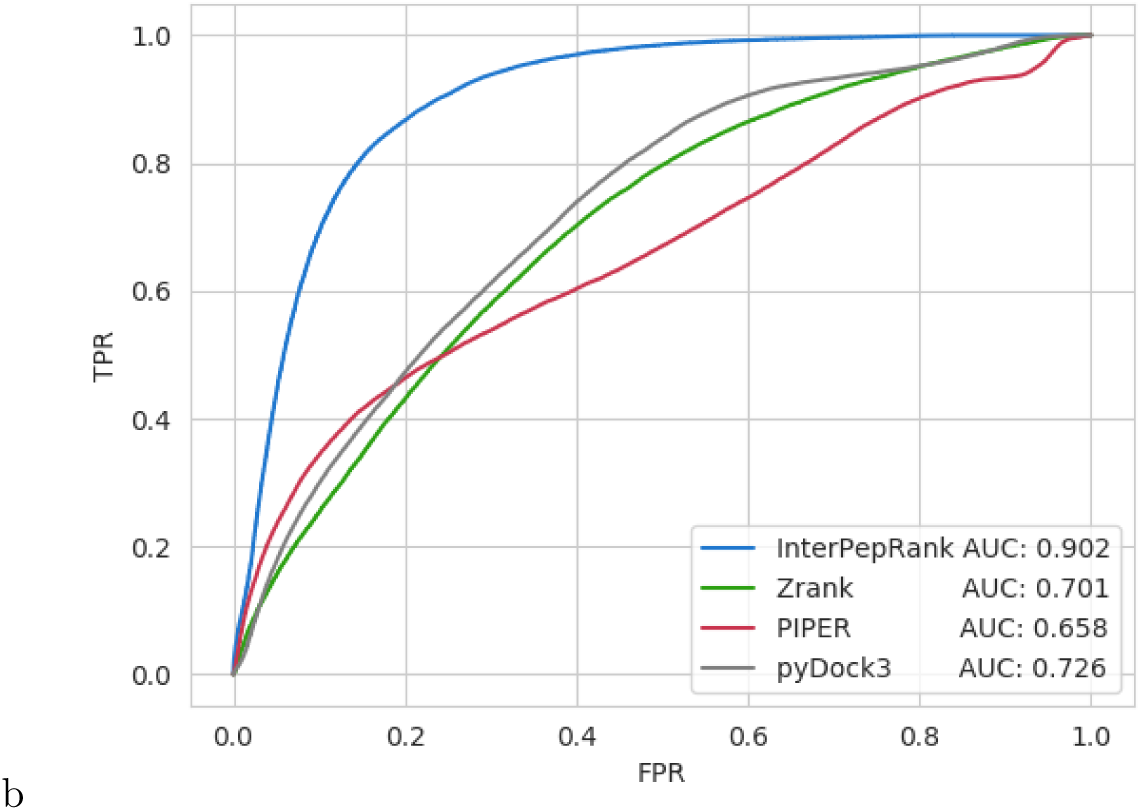
ROC-curves for using the method scores to separate low-LRMSD decoys from other decoys. Zrank, pyDock3, and PIPER are normalized by the total length of receptor and peptide. Analysis run only on the decoys of the Expanded Analysis set.

#### 3.1.1 Run-time and Large-scale Studies

Although slower than Zrank, InterPepRank still operates well within the realm of possibility of use in large-scale studies, see Table 2. Additionally, most of the time (circa 95%) of InterPepRank is spent preparing the graph-input to the net, something which could be optimized as compiled code rather than run as a python script.

**Table 2:** Average runtime over all targets for the different methods to evaluate 70,000 decoys, as measured in minutes, assuming all relevant files are available for each target. In the case of the 60 residue receptor and 472 residue receptor rows, it is instead the average runtime to evaluate 70,000 decoys over the 50 different peptide conformations. All calculations were performed on a single CPU core of Intel Xeon Gold 6130 running on CentOS 7, and when applicable an Nvidia GeForceRTX 2080 Ti graphics card was available. If InterPepRank is run completely on CPU, add approximately 30 minutes to the runtime.

| *Time (min)* | pyDock3 | Zrank | InterPepRank | Rosetta FPD scoring |
| --- | --- | --- | --- | --- |
| *Mean* | 664.4 | 9.2 | 100.6 | 590.0 |
| *Median* | 574.9 | 7.2 | 85.4 | 479.6 |
| *60 res. receptor* | 293.1 | 2.9 | 57.2 | 256.6 |
| *472 res. receptor* | 1372.6 | 24.7 | 249.2 | 1624.5 |

### 3.2 Proof of concept: Improving the PIPER-FlexPepDock pipeline

The purpose of InterPepRank is to provide an accurate rescoring step for selecting which decoys from FFT-based rigid-body-docking are worth further refining to achieve sub-Ångström docked complexes. The PIPER-FlexPepDock pipeline (Alam *et al.*, 2017) uses FFT-based PIPER to dock peptides of varying conformations onto a receptor surface, then utilizes PIPER-score to select 12,500 decoys for further refinement by the Rosetta FlexPepDock protocol, followed by clustering the top 1% of the refined decoys to make final docking predictions.

As a proof of concept, the full PIPER-FlexPepDock pipeline was run on the Expanded Analysis set both as is, and with InterPepRank, Zrank, or pyDock3 instead of PIPER for selecting decoys for refinement. The results were analyzed according to the CAPRI standard of classifying docked peptide-protein structures as compared to their native structures (Lensink *et al.*, 2017), and can be found in Table 3. It should be noted that many of the peptides in the Expanded Analysis set are longer than the maximum length investigated in the original PIPER-FlexPepDock paper, explaining the overall decrease in performance as compared to that paper. As seen in the table, both InterPepRank and Zrank are capable of improving the performance of PIPER-FlexPepDock if inserted as a function selecting decoys for refinement. While the inclusion of Zrank leads to a higher number of Acceptable models as per the CAPRI standard, the inclusion of InterPepRank leads to a higher yield of both Medium and High quality models. This is in line with the ROC-curves in Supplementary Figure 1 where Zrank can be seen to have fewer decoys for which it shows a ROC AUC below 0.5 than InterPepRank for this set, but where the average and median AUC of InterPepRank are higher.

**Table 3:** For how many of the targets in the Expanded Analysis set the modified PIPER-FlexPepDock pipeline produced models of the different quality measures among the top 10 results. As Rosetta FlexPepDock is a Monte-Carlo based approach, the part of the pipeline utilizing the FlexPepDock protocol and onwards was run in triplicates and the results averaged. Note that Acceptable means any model of at least Acceptable quality (same for Medium).

| *Re-scoring Method* | *Acceptable* | *Medium* | *High* |
| --- | --- | --- | --- |
| InterPepRank | 15.33 | **9.66** | **3.33** |
| Zrank | **18.0** | 7.0 | 2.33 |
| pyDock3 | 13.33 | 5.0 | 0.0 |
| None (PIPER) | 13.66 | 5.33 | 2.33 |

This is also reflected in the overall distribution LRMSD of the decoys sent for refinement by InterPepRank and Zrank: while InterPepRank both in mean and median selects more decoys with lower LRMSD, Zrank prefers a wider sampling of conformations resulting in a generally lower number of low LRMSD decoys, but meaning more targets have at least one low LRMSD decoy, see Supplementary Figures 1 and 2.

As InterPepRank has less targets with an AUC below 0.5 on the full test-set, we hypothesize that in the general case using InterPepRank to select decoys for refinement would be superior also in generating Acceptable models, as the larger full test set should be more representative of most real-world cases.

#### 3.2.1 Improving Run-time with Score Threshold

InterPepRank score is independent from protein and receptor size and composition, and only depends on the absolute quality of each decoy, which can be seen in Figure 4 showing InterPepRank can sort decoys even between targets. Because of this, it should be possible to introduce a score-threshold to avoid refining guaranteed bad decoys, as predicted by the InterPepRank score, rather than simply picking the top X decoys for refinement.

Indeed, adding an InterPepRank score cutoff of 0.47 for picking decoys for refinement rather than always picking the top 12,500 decoys reduces the median number of decoys refined per target to 10,000, while at least one High- or Medium-quality decoy will be produced for just as many targets as picking the top 12.500 decoys for refinement by InterPepRank without the threshold, and 95% of all targets which have at least one Acceptable decoy among the 12,500 will still have at least one.

Considering the median run-time of FlexPepDock refinement being 1.55 minutes per decoy on the same systems used for the re-scoring method speed benchmark, this results in the usage of an InterPepRank threshold saving on average 3,875 minutes of run-time, making up for the extra minutes spent running InterPepRank.

### 3.3 Example

As seen in previous works, machine learning methods for prediction of binding sites tend to predict protein-protein binding sites as potential peptide-protein binding sites, often resulting in difficulties in differentiating an interaction with the correct binding site from an interaction with another site, like one for protein-protein binding, or even crystal contacts (Johansson-Åkhe *et al.*, 2018, 2020a). A problem with shape-complementarity or electrostatics based methods on the other hand is their preference for conformations which maximize the number of contacts.

In this study, an example of a target difficult to both kinds of methods would be Siah1 (PDB ID: 4i7b). Its asymmetric unit, as well as many structural homologs, display it in a dimeric conformation, with the peptide binding at the opposite side of the protein. Additionally, there is a secondary hydrophobic groove which could maximize contacts with a potential peptide and has the same charge-distribution as the correct binding site, see Figure 5. While the true site shows an average evolutionary entropy of 0.520 and the dimerization site shows a similar entropy of 0.521, the hydrophobic groove shows less conservation with an entropy of 0.544 and the rest of the protein surface shows 0.603.

**Figure 5:**
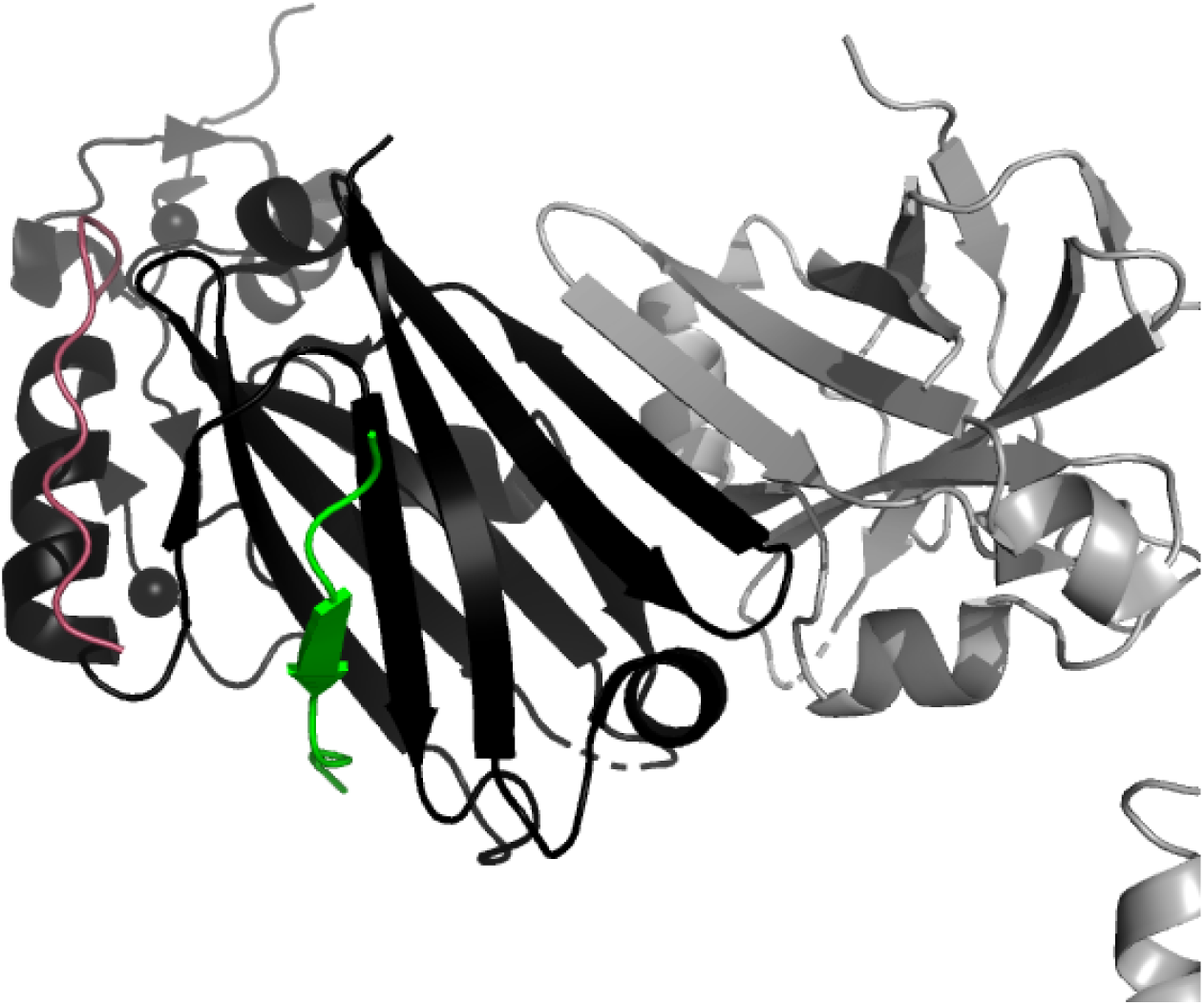
Siah1 in complex with a synthetic peptide (PDB-ID 4i7b). Siah1 is shown in black, the native conformation of the peptide is shown in green, the other chain in the asymmetric unit is shown in gray, and an example peptide conformation maximizing interaction in the alternate groove is shown in pink.

For this target, InterPepRank identified several locally favorable positions for the peptide, see Figure 6. The three wells roughly represent distances to the native peptide from decoys close to the native peptide, decoys bound at the hydrophobic groove by the helix, and decoys bound at the false dimerization site, respectively. These sites have average InterPepRank scores of 0.531, 0.496, and 0.465, respectively, while decoys bound over the rest of the protein surface average an InterPepRank score of 0.381. Similarly, the energy-based methods also report a lower average for the true binding site (−38.286 for Zrank and −3.150 for pyDock3) compared to the alternative sites (−30.4451 and −3.9 for the hydrophobic groove by the helix, and 4.145 and 3.108 for the dimerization site) and the rest of the protein surface (−7.142 and 4.742 respectively), albeit with some variation as pyDock3 generally prefers the groove by the helix to the correct site and Zrank ranks the dimerization site even worse than the rest of the surface. This demonstrates that InterPepRank does not simply select decoys arranged at conserved sites, but judges on more metrics, which is also supported by the fact that the length of the receptor multiple sequence alignment as well as the quality of alignments therein do not correlate with the quality of prediction (R ¡ 0.1), and that simpler Machine Learning models with the same architecture but less edge information perform worse (data not shown).

**Figure 6:**
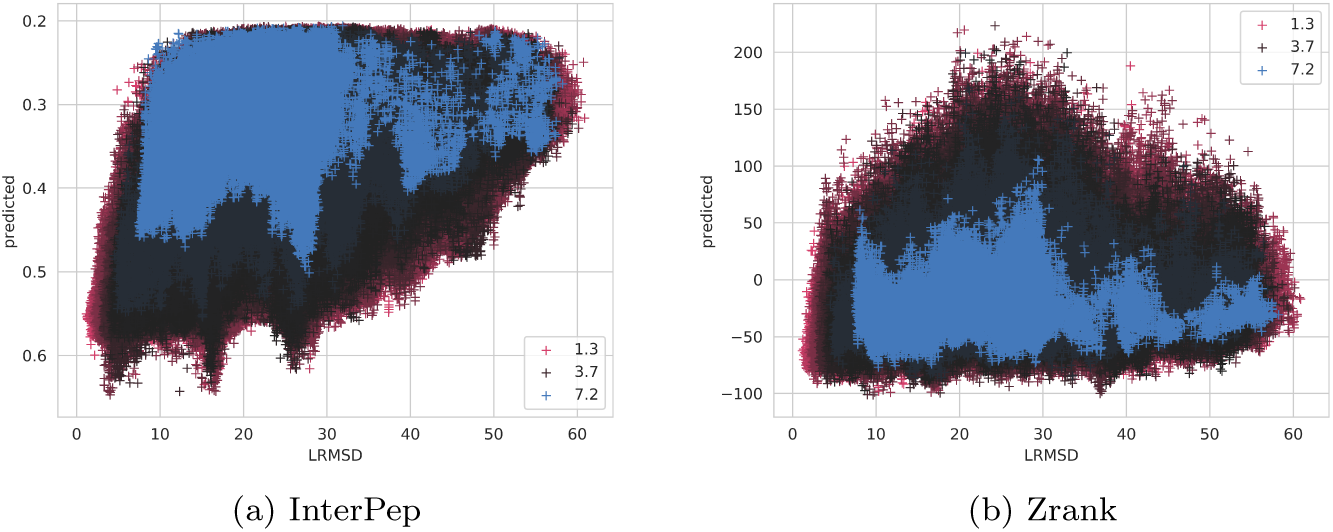
Scatterplot of InterPepRank predicted score (a) and Zrank predicted score (b) versus LRMSD for all decoys of 4i7b chains A (receptor) and B (peptide). Each decoy is colored by the backbone RMSD of the peptide to its native conformation if superimposed.

Similar scatter-plots for all test-targets and all methods benchmarked can be found in the Supplementary scatterplot compendium online (Johansson-Åkhe *et al.*, 2020b).

## 4 Conclusions

We have presented InterPepRank, a peptide-protein complex scoring and ranking method for use of rescoring and selecting coarse rigid-body-docking decoys for further refinement. InterPepRank uses both the structure of the complex and evolutionary features such as PSSM and sequence conservation to achieve high accuracy scoring in manageable computational time. The structure and the features are encoded in a graph representation where physical interactions between peptide and protein are represented as edges and the features are encoded in the nodes. The graph representation is trained using graph convolutions on a large set of peptide-protein complexes to predict the quality as measured by LRMSD. To maximize performance, the output of an ensemble of different network architectures are averaged in the final prediction.

On a massive independent test set not used to train and validate the method, InterPepRank has a median AUC of 0.86 for finding peptide-protein complexes with LRMSD¡4Å. This is an improvement compared to other ranking methods that have a median AUC of circa 0.65 to 0.69.

When inserted in the PIPER-FlexPepDock pipeline, InterPepRank consistently improves the performance of the pipeline by improving the selection of decoys for refinement, resulting in a 40% increase in High quality models produced, and a 80% increase in Medium quality models produced, with the possibility of not increasing the overall computational time without loss in performance by filtering decoys by InterPepRank score.

In addition to selecting peptide-protein complexes for all-atom refinement, InterPepRank should prove useful for providing a cross-target comparable scoring function.

## Acknowledgments

This work was supported by a Swedish Research Council grant, 2016-05369, The Swedish e-Science Research Center, and the Foundation Blanceflor Boncompagni Ludovisi, née Bildt. The computations were performed on resources provided by the Swedish National Infrastructure for Computing (SNIC) at the National Supercomputer Centre (NSC) in Linköping.

## Notes

https://doi.org/10.17044/scilifelab.13134756

### Competing Interest Statement

The authors have declared no competing interest.

### Summary of Updates

Test-set greatly expanded, affecting Figures 3 and 4, with minor additions to Results and Discussion as necessary. Sections throughout the paper, especially regarding Methods, have been clarified.

