## Supplementary for "InterPepRank: Assessment of Docked Peptide Conformations by a Deep Graph Network"

### InterPepRank Supplementary Information

#### 1 Architecture

In Tables 1, 2, 3, 4, 5, 9, and 10, the detailed architectures for the 9 network architectures considered in the final ensemble for InterPepDeep can be found. The nets are slight variations on the same basic architecture, as described in the main paper.

| **Name** | **Layer** | **Dimensions** | **Input** |
|---|---|---|---|
| Node Cons. Features | input | 100×42 | - |
| Node Ligand Var. | input | 100×1 | - |
| Amino Acid One-hot | input | 100×21 | - |
| Edge Features Input | input | 100×100×4 | - |
| Amino Acid Embed | embedding | 100×4 | Amino Acid One-hot |
| Node Features | concatenate | 100×47 | Amino Acid Embed<br>Node Cons. Features<br>Node Ligand Var. |
| Edge Features | 2D dropout(25%) | 100×100×4 | Edge Features Input |
| EdgeConv1 | edge conditioned convolution<br>(ReLU activation) | 100×8 | Edge Features<br>Node Features |
| EdgeConv2 | edge conditioned convolution<br>(ReLU activation) | 100×8 | Edge Features Input<br>EdgeConv1 |
| EdgeConv3 | edge conditioned convolution<br>(ReLU activation) | 100×16 | Edge Features Input<br>EdgeConv2 |
| EdgeConv4 | edge conditioned convolution<br>(ReLU activation) | 100×16 | Edge Features Input<br>EdgeConv3 |
| Concatenate | concatenate | 100×48 | EdgeConv1<br>EdgeConv2<br>EdgeConv3<br>EdgeConv4 |
| Pooling | GlobalAveragePooling | 48 | Concatenate |
| Dropout | dropout (25%) | 48 | Pooling |
| Dense | dense | 32 | Dropout |
| Activation | ReLU activation | 32 | Dense |
| Classifier | dense | 4 | Activation |
| Output | softmax | 4 | Classifier |

Tab. 1: Architecture for net 0 considered in the ensemble-prediction of InterPepDeep. The 4 classes are evenly distributed over the range 0 to 1 as the net predicts the S-score normalized LRMSD (normalized with 4.0 LRMSD as 0.5).

| Name | Layer | Dimensions | Input |
|---|---|---|---|
| Node Cons. Features | input | 100×42 | - |
| Node Ligand Var. | input | 100×1 | - |
| Amino Acid One-hot | input | 100×21 | - |
| Edge Features Input | input | 100×100×4 | - |
| Amino Acid Embed | embedding | 100×4 | Amino Acid One-hot |
| Node Features | concatenate | 100×47 | Amino Acid Embed<br>Node Cons. Features<br>Node Ligand Var. |
| Edge Features | 2D dropout(25%) | 100×100×4 | Edge Features Input |
| EdgeConv1 | edge conditioned convolution<br>(ReLU activation) | 100×8 | Edge Features<br>Node Features |
| EdgeConv2 | edge conditioned convolution<br>(ReLU activation) | 100×8 | Edge Features Input<br>EdgeConv1 |
| EdgeConv3 | edge conditioned convolution<br>(ReLU activation) | 100×16 | Edge Features Input<br>EdgeConv2 |
| EdgeConv4 | edge conditioned convolution<br>(ReLU activation) | 100×16 | Edge Features Input<br>EdgeConv3 |
| Concatenate | concatenate | 100×48 | EdgeConv1<br>EdgeConv2<br>EdgeConv3<br>EdgeConv4 |
| Pooling | GlobalAveragePooling | 48 | Concatenate |
| Dropout | dropout (25%) | 48 | Pooling |
| Dense | dense | 16 | Dropout |
| Activation | ReLU activation | 16 | Dense |
| Classifier | dense | 3 | Activation |
| Output | softmax | 3 | Classifier |

Tab. 2: Architecture for net 1 considered in the ensemble-prediction of InterPepRank. The 3 classes were the interval from 0.0 to 1.0 segmented by 0.75 and 0.5 as the net predicts the S-score normalized LRMSD (normalized with 4.0 LRMSD as 0.5).

| Name | Layer | Dimensions | Input |
|---|---|---|---|
| Node Cons. Features | input | 100×42 | - |
| Node Ligand Var. | input | 100×1 | - |
| Amino Acid One-hot | input | 100×21 | - |
| Edge Features Input | input | 100×100×4 | - |
| Amino Acid Embed | embedding | 100×4 | Amino Acid One-hot |
| Node Features | concatenate | 100×47 | Amino Acid Embed<br>Node Cons. Features<br>Node Ligand Var. |
| Edge Features | 2D dropout(25%) | 100×100×4 | Edge Features Input |
| EdgeConv1 | edge conditioned convolution<br>(ReLU activation) | 100×8 | Edge Features<br>Node Features |
| EdgeConv2 | edge conditioned convolution<br>(ReLU activation) | 100×8 | Edge Features Input<br>EdgeConv1 |
| EdgeConv3 | edge conditioned convolution<br>(ReLU activation) | 100×16 | Edge Features Input<br>EdgeConv2 |
| EdgeConv4 | edge conditioned convolution<br>(ReLU activation) | 100×16 | Edge Features Input<br>EdgeConv3 |
| Concatenate | concatenate | 100×48 | EdgeConv1<br>EdgeConv2<br>EdgeConv3<br>EdgeConv4 |
| Pooling | GlobalAveragePooling | 48 | Concatenate |
| Dropout | dropout (25%) | 48 | Pooling |
| Dense | dense | 16 | Dropout |
| Activation | ReLU activation | 16 | Dense |
| Classifier | dense | 2 | Activation |
| Output | softmax | 2 | Classifier |

Tab. 3: Architecture for net 2 considered in the ensemble-prediction of InterPepRank.

| **Name** | **Layer** | **Dimensions** | **Input** |
|---|---|---|---|
| Node Cons. Features | input | 100×42 | - |
| Node Ligand Var. | input | 100×1 | - |
| Amino Acid One-hot | input | 100×21 | - |
| Edge Features Input | input | 100×100×4 | - |
| Amino Acid Embed | embedding | 100×2 | Amino Acid One-hot |
| Node Features | concatenate | 100×45 | Amino Acid Embed<br>Node Cons. Features<br>Node Ligand Var. |
| Edge Features | 2D dropout(25%) | 100×100×4 | Edge Features Input |
| EdgeConv1 | edge conditioned convolution<br>(ReLU activation) | 100×8 | Edge Features<br>Node Features |
| EdgeConv2 | edge conditioned convolution<br>(ReLU activation) | 100×8 | Edge Features Input<br>EdgeConv1 |
| EdgeConv3 | edge conditioned convolution<br>(ReLU activation) | 100×16 | Edge Features Input<br>EdgeConv2 |
| EdgeConv4 | edge conditioned convolution<br>(ReLU activation) | 100×16 | Edge Features Input<br>EdgeConv3 |
| Concatenate | concatenate | 100×48 | EdgeConv1<br>EdgeConv2<br>EdgeConv3<br>EdgeConv4 |
| Pooling | GlobalAveragePooling | 48 | Concatenate |
| Dropout | dropout (25%) | 48 | Pooling |
| Dense | dense | 32 | Dropout |
| Activation | ReLU activation | 32 | Dense |
| Classifier | dense | 2 | Activation |
| Output | softmax | 2 | Classifier |

Tab. 4: Architecture for net 3 considered in the ensemble-prediction of InterPepRank.

| Name | Layer | Dimensions | Input |
|---|---|---|---|
| Node Cons. Features | input | 100×42 | - |
| Node Ligand Var. | input | 100×1 | - |
| Amino Acid One-hot | input | 100×21 | - |
| Edge Features Input | input | 100×100×4 | - |
| Amino Acid Embed | embedding | 100×4 | Amino Acid One-hot |
| Node Features | concatenate | 100×47 | Amino Acid Embed<br>Node Cons. Features<br>Node Ligand Var. |
| Edge Features | 2D dropout(10%) | 100×100×4 | Edge Features Input |
| EdgeConv1 | edge conditioned convolution<br>(kernel net 8, ReLU activation) | 100×8 | Edge Features<br>Node Features |
| EdgeConv2 | edge conditioned convolution<br>(kernel net 8, ReLU activation) | 100×8 | Edge Features Input<br>EdgeConv1 |
| EdgeConv3 | edge conditioned convolution<br>(kernel net 16, ReLU activation) | 100×16 | Edge Features Input<br>EdgeConv2 |
| EdgeConv4 | edge conditioned convolution<br>(kernel net 16, ReLU activation) | 100×16 | Edge Features Input<br>EdgeConv3 |
| Concatenate | concatenate | 100×48 | EdgeConv1<br>EdgeConv2<br>EdgeConv3<br>EdgeConv4 |
| Pooling | GlobalAveragePooling | 48 | Concatenate |
| Dropout | dropout (10%) | 48 | Pooling |
| Dense | dense | 32 | Dropout |
| Activation | ReLU activation | 32 | Dense |
| Classifier | dense | 2 | Activation |
| Output | softmax | 2 | Classifier |

Tab. 5: Architecture for net 4 considered in the ensemble-prediction of InterPepRank.

| Name | Layer | Dimensions | Input |
|---|---|---|---|
| Node Cons. Features | input | 50×42 | - |
| Node Ligand Var. | input | 50×1 | - |
| Amino Acid One-hot | input | 50×21 | - |
| Edge Features Input | input | 50×50×4 | - |
| Amino Acid Embed | embedding | 50×4 | Amino Acid One-hot |
| Node Features | concatenate | 50×47 | Amino Acid Embed<br>Node Cons. Features<br>Node Ligand Var. |
| Edge Features | 2D dropout(10%) | 50×50×4 | Edge Features Input |
| EdgeConv1 | edge conditioned convolution<br>(ReLU activation) | 50×8 | Edge Features<br>Node Features |
| EdgeConv2 | edge conditioned convolution<br>(ReLU activation) | 50×8 | Edge Features Input<br>EdgeConv1 |
| EdgeConv3 | edge conditioned convolution<br>(ReLU activation) | 50×16 | Edge Features Input<br>EdgeConv2 |
| EdgeConv4 | edge conditioned convolution<br>(ReLU activation) | 50×16 | Edge Features Input<br>EdgeConv3 |
| Concatenate | concatenate | 50×48 | EdgeConv1<br>EdgeConv2<br>EdgeConv3<br>EdgeConv4 |
| Pooling | GlobalAveragePooling | 48 | Concatenate |
| Dropout | dropout (10%) | 48 | Pooling |
| Dense | dense | 32 | Dropout |
| Activation | ReLU activation | 32 | Dense |
| Classifier | dense | 2 | Activation |
| Output | softmax | 2 | Classifier |

Tab. 6: Architecture for net 5 considered in the ensemble-prediction of InterPepRank. The input for this net was constructed the same way as for the other networks, but with a limit of 50 residues rather than 100.

| Name | Layer | Dimensions | Input |
|---|---|---|---|
| Node Cons. Features | input | 100×42 | - |
| Node Ligand Var. | input | 100×1 | - |
| Amino Acid One-hot | input | 100×21 | - |
| Node Features | concatenate | 100×64 | Node Cons. Features<br>Node Ligand Var.<br>Amino Acid One-hot |
| Edge Features | input | 100×100×4 | - |
| EdgeConv1 | edge conditioned convolution<br>(kernel net 8, ReLU activation) | 100×8 | Edge Features<br>Node Features |
| BatchNorm1 | batch normalization | 100×8 | EdgeConv1 |
| Dropout1 | dropout (10%) | 100×8 | BatchNorm1 |
| EdgeConv2 | edge conditioned convolution<br>(kernel net 8, ReLU activation) | 100×8 | Edge Features<br>Dropout1 |
| BatchNorm2 | batch normalization | 100×8 | EdgeConv2 |
| Dropout2 | dropout (10%) | 100×8 | BatchNorm2 |
| Bypass1 | 1D dense | 100×8 | Node Features |
| Block1 | addition | 100×8 | Dropout2<br>Bypass1 |
| EdgeConv3 | edge conditioned convolution<br>(kernel net 8, ReLU activation) | 100×8 | Edge Features<br>Block1 |
| BatchNorm3 | batch normalization | 100×8 | EdgeConv3 |
| Dropout3 | dropout (10%) | 100×8 | BatchNorm3 |
| EdgeConv4 | edge conditioned convolution<br>(kernel net 8, ReLU activation) | 100×8 | Edge Features<br>Dropout3 |
| BatchNorm4 | batch normalization | 100×8 | EdgeConv4 |
| Dropout4 | dropout (10%) | 100×8 | BatchNorm4 |
| Block2 | addition | 100×8 | Dropout4<br>Block1 |
| EdgeConv5 | edge conditioned convolution<br>(kernel net 8, ReLU activation) | 100×8 | Edge Features<br>Block2 |
| BatchNorm5 | batch normalization | 100×8 | EdgeConv5 |
| Dropout5 | dropout (10%) | 100×8 | BatchNorm5 |
| EdgeConv6 | edge conditioned convolution<br>(kernel net 8, ReLU activation) | 100×8 | Edge Features<br>Dropout5 |
| BatchNorm6 | batch normalization | 100×8 | EdgeConv6 |
| Dropout6 | dropout (10%) | 100×8 | BatchNorm6 |
| Block3 | addition | 100×8 | Dropout6<br>Block2 |
| EdgeConv7 | edge conditioned convolution<br>(kernel net 8, ReLU activation) | 100×8 | Edge Features<br>Block3 |
| BatchNorm7 | batch normalization | 100×8 | EdgeConv7 |
| Dropout7 | dropout (10%) | 100×8 | BatchNorm7 |
| EdgeConv8 | edge conditioned convolution<br>(kernel net 8, ReLU activation) | 100×8 | Edge Features<br>Dropout7 |
| BatchNorm8 | batch normalization | 100×8 | EdgeConv8 |
| Dropout8 | dropout (10%) | 100×8 | BatchNorm8 |
| Block4 | addition | 100×8 | Dropout8<br>Block3 |
| EdgeConv9 | edge conditioned convolution<br>(kernel net 8, ReLU activation) | 100×32 | Edge Features<br>Block4 |
| Pooling | GlobalAttentionPool | 32 | EdgeConv9 |
| BatchNorm9 | batch normalization | 32 | EdgeConv9 |
| Dropout9 | dropout (10%) | 32 | BatchNorm9 |
| Prediction | dense | 1 | Dropout9 |

Tab. 7: Architecture for net 6 considered in the ensemble-prediction of InterPepRank. Training of net 6 was done in a binary connected manner, running two copies of the net in parallel with weight-sharing inbetween on two different decoys at any given moment. Additionally, during training another net found in Table 8 was attached to the binary net, and the loss function was calculated on this net's capacity to classify which of the two decoys is closer to native, as well as the individual losses from the single branches, weighting single branches 0.1 and the comparison at 1.0. This approach is similar to the Tricephalous net suggested by Hurtado *et al.* (2018).

| Name | Layer | Dimensions | Input |
| --- | --- | --- | --- |
| EdgeConv10 | edge conditioned convolution (kernel net 16, ReLU activation) weight-sharing with EdgeConv11 | 100×16 | Edge Features 6_1 Block4 6_1 |
| EdgeConv11 | edge conditioned convolution (kernel net 16, ReLU activation) weight-sharing with EdgeConv10 | 100×16 | Edge Features 6_2 Block4 6_2 |
| EdgeConv12 | edge conditioned convolution (kernel net 16, ReLU activation) weight-sharing with EdgeConv13 | 100×16 | Edge Features 6_1 Block4 6_1 |
| EdgeConv13 | edge conditioned convolution (kernel net 16, ReLU activation) weight-sharing with EdgeConv12 | 100×16 | Edge Features 6_2 Block4 6_2 |
| Pooling1 | GlobalAttentionPool (32) | 32 | EdgeConv10 |
| Pooling2 | GlobalAttentionPool (32) | 32 | EdgeConv11 |
| Pooling3 | GlobalAttentionPool (32) | 32 | EdgeConv12 |
| Pooling4 | GlobalAttentionPool (32) | 32 | EdgeConv13 |
| Add1 | addition | 32 | Pooling1 Pooling4 |
| Add2 | addition | 32 | Pooling2 Pooling3 |
| BatchNorm10 | batch normalization | 32 | Add1 |
| Dropout10 | dropout (10%) | 32 | BatchNorm10 |
| BatchNorm11 | batch normalization | 32 | Add2 |
| Dropout11 | dropout (10%) | 32 | BatchNorm11 |
| Dense1 | dense | 1 | Dropout10 |
| Dense2 | dense | 1 | Dropout11 |
| Comparison | addition | 1 | Dense1 Dense2 |

Tab. 8: The extra comparison-net used during training of net 6, attached to two weight-sharing instances of net 6 referred to as net 6_1 and 6_2.

| Name | Layer | Dimensions | Input |
|---|---|---|---|
| Node Cons. Features | input | 100×42 | - |
| Node Ligand Var. | input | 100×1 | - |
| Amino Acid One-hot | input | 100×21 | - |
| Edge Features Input | input | 100×100×4 | - |
| Amino Acid Embed | embedding | 100×4 | Amino Acid One-hot |
| Node Features | concatenate | 100×47 | Amino Acid Embed<br>Node Cons. Features<br>Node Ligand Var. |
| Edge Features | 2D dropout(25%) | 100×100×4 | Edge Features Input |
| EdgeConv1 | edge conditioned convolution<br>(kernel net 8, ReLU activation) | 100×8 | Edge Features<br>Node Features |
| EdgeConv2 | edge conditioned convolution<br>(kernel net 8, ReLU activation) | 100×8 | Edge Features<br>EdgeConv1 |
| EdgeConv3 | edge conditioned convolution<br>(kernel net 16, ReLU activation) | 100×16 | Edge Features<br>EdgeConv2 |
| EdgeConv4 | edge conditioned convolution<br>(kernel net 16, ReLU activation) | 100×16 | Edge Features<br>EdgeConv3 |
| Concatenate | concatenate | 100×48 | EdgeConv1<br>EdgeConv2<br>EdgeConv3<br>EdgeConv4 |
| Pooling | GlobalAttentionPooling | 32 | Concatenate |
| Dropout | dropout (25%) | 32 | Pooling |
| Dense | dense | 32 | Dropout |
| Activation | ReLU activation | 32 | Dense |
| Classifier | dense | 4 | Activation |
| Output | softmax | 4 | Classifier |

Tab. 9: Architecture for net 7 considered in the ensemble-prediction of InterPepRank. The 4 classes are evenly distributed over the range 0 to 1 as the net predicts the S-score normalized LRMSD (normalized with 4.0 LRMSD as 0.5).

| Name | Layer | Dimensions | Input |
|---|---|---|---|
| Node Cons. Features | input | 100×42 | - |
| Node Ligand Var. | input | 100×1 | - |
| Amino Acid One-hot | input | 100×21 | - |
| Edge Features Input | input | 100×100×4 | - |
| Amino Acid Embed | embedding | 100×4 | Amino Acid One-hot |
| Node Features | concatenate | 100×47 | Amino Acid Embed<br>Node Cons. Features<br>Node Ligand Var. |
| Edge Features | 2D dropout(10%) | 100×100×4 | Edge Features Input |
| EdgeConv1 | edge conditioned convolution<br>(kernel net 8, ReLU activation) | 100×8 | Edge Features<br>Node Features |
| EdgeConv2 | edge conditioned convolution<br>(kernel net 8, ReLU activation) | 100×8 | Edge Features<br>EdgeConv1 |
| EdgeConv3 | edge conditioned convolution<br>(kernel net 16, ReLU activation) | 100×16 | Edge Features<br>EdgeConv2 |
| EdgeConv4 | edge conditioned convolution<br>(kernel net 16, ReLU activation) | 100×16 | Edge Features<br>EdgeConv3 |
| Concatenate | concatenate | 100×48 | EdgeConv1<br>EdgeConv2<br>EdgeConv3<br>EdgeConv4 |
| Pooling | GlobalAttentionPooling | 32 | Concatenate |
| Dropout | dropout (10%) | 32 | Pooling |
| Dense | dense | 32 | Dropout |
| Activation | ReLU activation | 32 | Dense |
| Classifier | dense | 2 | Activation |
| Output | softmax | 2 | Classifier |

Tab. 10: Architecture for net 8 considered in the ensemble-prediction of InterPepRank.

#### 2 Expanded Analysis test set

With a decrease in test set size, computationally heavy re-scoring methods like pyDock3 or Rosetta FlexPepDock scoring-mode can be included in the comparison. See Figure 1 for an analogue to Figure 4 of the main paper.

Using Rosetta FlexPepDock scoring mode only to re-score rigid-body docked decoys proved slow, even without any refinement, as was discussed in the main paper. Since the Rosetta scoring function is a fine-grained function developed for protein refinement and design, it makes sense it would perform poorly on structures not necessarily optimal by the Rosetta standard. Indeed, when using Rosetta to score structures, it is common practice to first relax the structure through the Rosetta Relax protocol, something which would considerably add to the run-time if attempted in this situation.

#### 3 Decoy Distribution

In Figure 2 are some graphical representations of the LRMSD distributions of decoys selected by the different scoring methods for refinement. Results are only shown for the set all methods were run on.

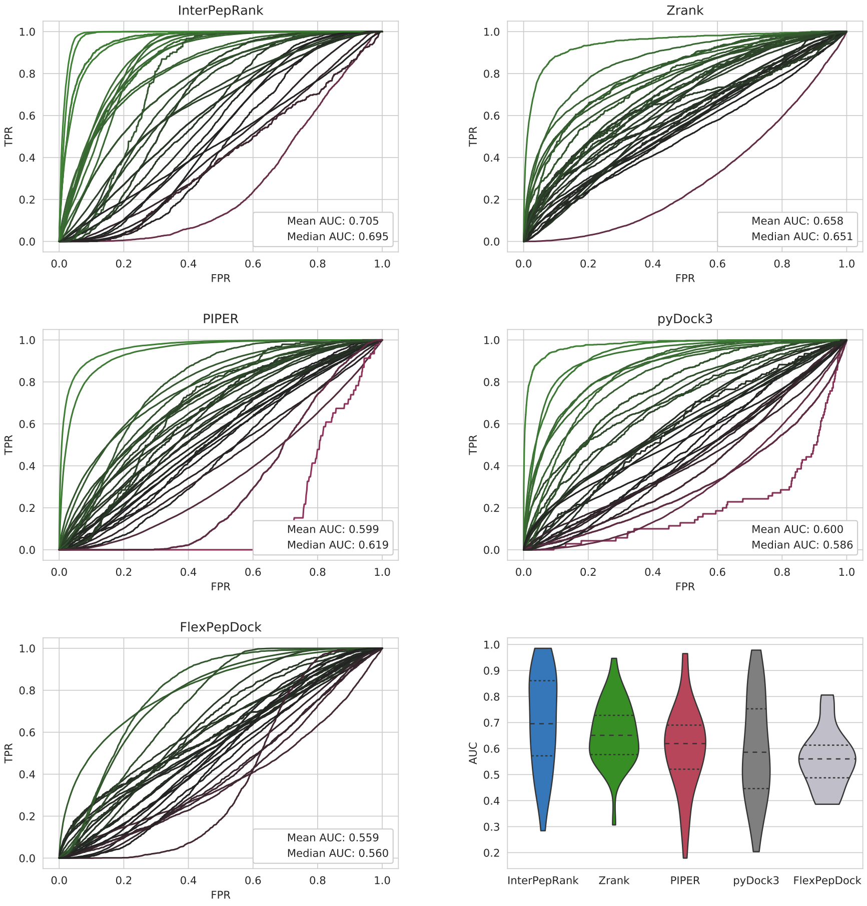


Fig. 1: ROC-curves for all methods discussed in the main paper, including pyDock3 and Rosetta FlexPepDock scoring mode and a violin-curve summarizing all AUCs, for the Expanded Analysis set (a randomly selected set of 50 targets all methods were run on). Each target is represented by 1 curve. The area under the curve (AUC) displayed in the graphs is the average and median over all targets.

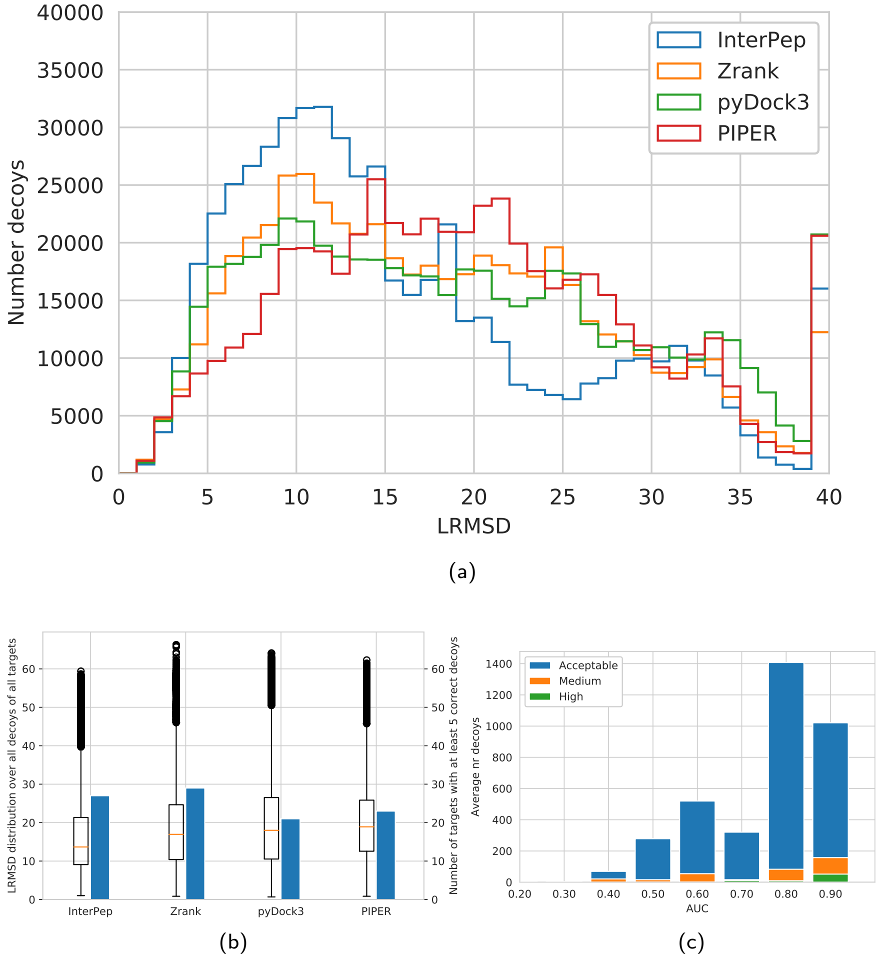


Fig. 2: Distribution of LRMSD of decoys selected for refinement by the different methods. Results shown for the Expanded Analysis set. In (a), all decoys at LRMSD 40+ were summed into the 40 Å bin. In (c), the median number of models of the different quality-measures produced per method per target after refinement for all methods and targets in the Expanded Analysis set are shown binned by AUC on original decoys, to highlight that good performance on the rigid-body docked decoys translates to well-refined models.
